# Transcriptome of epibiont Saccharibacteria *Nanosynbacter lyticus* strain TM7x during establishment of symbiosis

**DOI:** 10.1101/2022.03.25.485537

**Authors:** Erik L Hendrickson, Batbileg Bor, Kristopher A. Kerns, Eleanor I. Lamont, Yunjie Chang, Jun Liu, Lujia Cen, Wenyuan Shi, Xuesong He, Jeffrey S McLean

## Abstract

Saccharibacteria *Nanosynbacter lyticus* strain TM7x is a member of the broadly distributed Candidate Phylum Radiation. These bacteria have ultrasmall cell size, reduced genomes and live as epibionts on the surface of other bacteria. The mechanisms by which they establish and maintain this relationship are not yet fully understood. The transcriptomes of the epibiont TM7x and its host bacteria *Schaalia odontolyticus* strain XH001 were captured across the establishment of symbiosis during both the initial interaction and stable symbiosis. The results showed a dynamic interaction with large shifts in gene expression for both species between the initial encounter and stable symbiosis, notably transporter genes. During stable symbiosis, the host XH001 showed higher gene expression for peptidoglycan biosynthesis, mannosylation, cell cycle and stress related genes, but lower expression of chromosomal partitioning genes. This was consistent with the elongated cell shape seen in XH001 infected with TM7x and our discovery that infection resulted in thickened cell walls. Within TM7x, increased pili, type IV effector gene, and arginine catabolism/biosynthesis gene expression during stable symbiosis implied a key role for these functions in the interaction. Consistent with its survival and persistence in the human microbiome as an obligate epibiont with reduced *de novo* biosynthetic capacities, TM7x also showed higher levels for energy production and peptidoglycan biosynthesis but lower expression of stress related genes during stable symbiosis. These results imply that TM7x and its host bacteria keep a delicate balance in order to sustain an episymbiotic lifestyle.

**IMPORTANCE:** *Nanosynbacter lyticus* type strain TM7x is the first cultivated member of the Saccharibacteria and the Candidate Phyla Radiation (CPR). It was discovered to have ultrasmall cell size with a highly reduced genome that establishes an obligate epibiotic relationship with its host bacterium. The CPR, now formally proposed as the Patescibacteria super-phylum, is a large monophyletic radiation of diverse bacteria with reduced genomes that includes Saccharibacteria. The vast majority of the CPR have yet to be cultivated in the laboratory and our insights into these unique organisms to date has been derived from only a few Saccharibacteria species. It is unknown however how these small obligate parasitic Saccharibacteria, that are missing many *de novo* biosynthetic pathways, are maintained at high prevalence within the human microbiome as well as in the environment. When TM7x infects its host bacterium there are distinct temporal phases, including an initial interaction, a killing phase, recovery phase, and finally stable symbiosis. Here we captured the gene expression of the host bacterium and epibiont during this dynamic interaction which represents the initial insights into the mechanisms of how these unique microbes may survive and persist.

## INTRODUCTION

Saccharibacteria *Nanosynbacter lyticus* type strain TM7x has an ultrasmall cell size (200- 300nm) and a highly reduced genome. It was originally isolated from the human oral cavity as an epibiont on a bacterial host (1), *Schaalia odontolyticus* strain XH001 (formerly *Actinomyces odontolyticus*) (2). *N. lyticus* strain TM7x later became the first cultivated species of a large bacterial group discovered from filtered groundwater denoted as the Candidate Phyla Radiation (CPR), a monophyletic group with reduced genomes, also now referred to as Patescibacteria super- phylum (1, 3–5). Genomes from the yet-to-be-cultivated CPR members are also noted for reduced genomes with minimal biosynthetic capabilities (3, 4, 6). Recent work has demonstrated high prevalence and a large genomic diversity of Saccharibacteria in the human microbiome, and across mammals (5, 7, 8). The currently recognized broad presence of these bacterial epibionts generates much interest in understanding the mechanisms underlying their obligate symbiosis. Specifically, how do these ultrasmall bacteria with limited biosynthetic capacity survive and persist in these very diverse niches.

Despite being obligate epibionts with reduced *de novo* biosynthetic capacity, the mammalian associated Saccharibacteria have remarkable genomic diversity. One of the six major groups (G1-G6) (5, 9), the G1, uniquely consists of both mammalian members and environmental representatives, sharing ∼60% of their protein coding genes (5). Furthermore, *N. lyticus* strain TM7x and other recently isolated G1 strains from the oral cavity have retained a highly syntenic gene arrangement with environmental counterparts (1, 5, 7, 10). All oral Saccharibacteria genomes appear to have undergone further reductive evolution during the adaptation and evolution in mammals. As with other members of the CPR, TM7x is missing a number of major pathways including tricarboxylic acid cycle and *de novo* synthesis of nucleotides and amino acids. However, of the recent fully closed genomes of sequenced isolates (10, 11) from the G1 group, TM7x has the most streamlined genome. Initial analysis of the TM7x genome predicted 739 genes (1), 692 protein encoding, densely coded with a percentage coding base count of 94%. This results in short intergenic regions with no currently validated small RNAs, although only one global transcriptome study at a single timepoint has been conducted to date (1).

Through extensive *in vitro* studies on the dynamics of the XH001/TM7x interaction, it was found that while unable to grow without a host bacterium, free-floating TM7x could be isolated from a coculture and infect a new host horizontally (12). It was also observed that naive bacterial hosts, those that were never part of an established coculture, underwent an unusual pattern during passaging. First was a killing phase where host numbers decreased dramatically while large numbers of TM7x infected single host cells (12). This was followed by a recovery phase where host numbers returned to original levels while the number of TM7x on each host cell decreased. Finally, after a certain number of passages, the coculture reached a stable symbiosis between TM7x and the host XH001 and no longer showed drastic killing. Explorations of the TM7x-host range found that the epibiont was able to consistently grow on a single group of phylogenetically related *Schaalia* species, revealing this strain’s restricted host-range (13). While there was variation when tested on multiple TM7x host bacteria, most TM7x infections followed the same pattern as XH001; initial encounter, killing phase, recovery phase, and finally stable symbiosis.

To better understand the dynamics between XH001/TM7x across these different phases towards the establishment of stable symbiosis, we investigated the temporal transcriptional profiles of both species after initiation of infection. This work characterizes the initial encounter as well as the temporal expression dynamics for a member of the CPR. Extensive expression changes were found between the initial encounter and stable symbiosis in both XH001 and TM7x, including changes in energy metabolism, transporters, and cell wall metabolism, enabling further insights into the lifestyle of this unique, mammalian-associated, ultrasmall epibiont with a highly reduced genome.

## MATERIALS AND METHODS

### Bacterial Strains and Growth Conditions

Naive XH001 (XH001n) (*Schaalia odontolyticus* subsp. *Actinosynbacter* strain XH001) monoculture and XH001/TM7x coculture were both grown in Bacto Brain Heart Infusion (BHI, BD) medium at 37 °C in a microaerophilic chamber (2% O2, 5% CO2, balanced with N2) (12, 13). Free floating TM7x (*Nanosynbacter lyticus* Strain TM7x HMT-952) was isolated from XH001/TM7x coculture as described previously (12). Briefly, XH001/TM7x coculture were filtered through 0.45 um filter and TM7x cells were collected by ultracentrifugation at 80,000 x g for 90 min. Isolated TM7x cells were used freshly on the same day for infection experiments without freezing or overnight storage.

### TM7x Infection, Passaging and Sample Collection

Three independent cultures of XH001n cells were recovered from frozen stock and passaged twice in BHI every 24 hours to ensure homogeneity before used for setting up TM7x infection assay as previously described with modification (12). Briefly, 115 mL of 0.1 OD600 XH001n cells were mixed at a roughly 1:1 ratio with isolated free floating TM7x cells in triplicate (12). After 6 hours of incubation, the first time point (Passage 0) was sampled from each replicate (passage 0, hour 6). 50 mL of culture were centrifuged at 5,500 rpm for 10 minutes at 4°C. Supernatants were discarded, and the pellets were flash-frozen in liquid nitrogen. Remaining cultures were grown to the 24-hour mark and XH001 cell density measurement by OD600 and CFU by plating were determined at each passage. At each passage, 10-50 mL cultures were diluted to 0.1 OD600. The volume depended on the follow up collection and sampling. In the sixth passage, after 6 hours (passage 6, hour 6), samples of 50 mL culture were collected, pelleted, and flash-frozen. We also conducted a parallel experiment where XH001n alone, without the addition of TM7x, underwent the same inoculation, passaging and sampling. Both OD600 and CFU were recorded and graphed in **Figure 1**.

**Figure 1.**
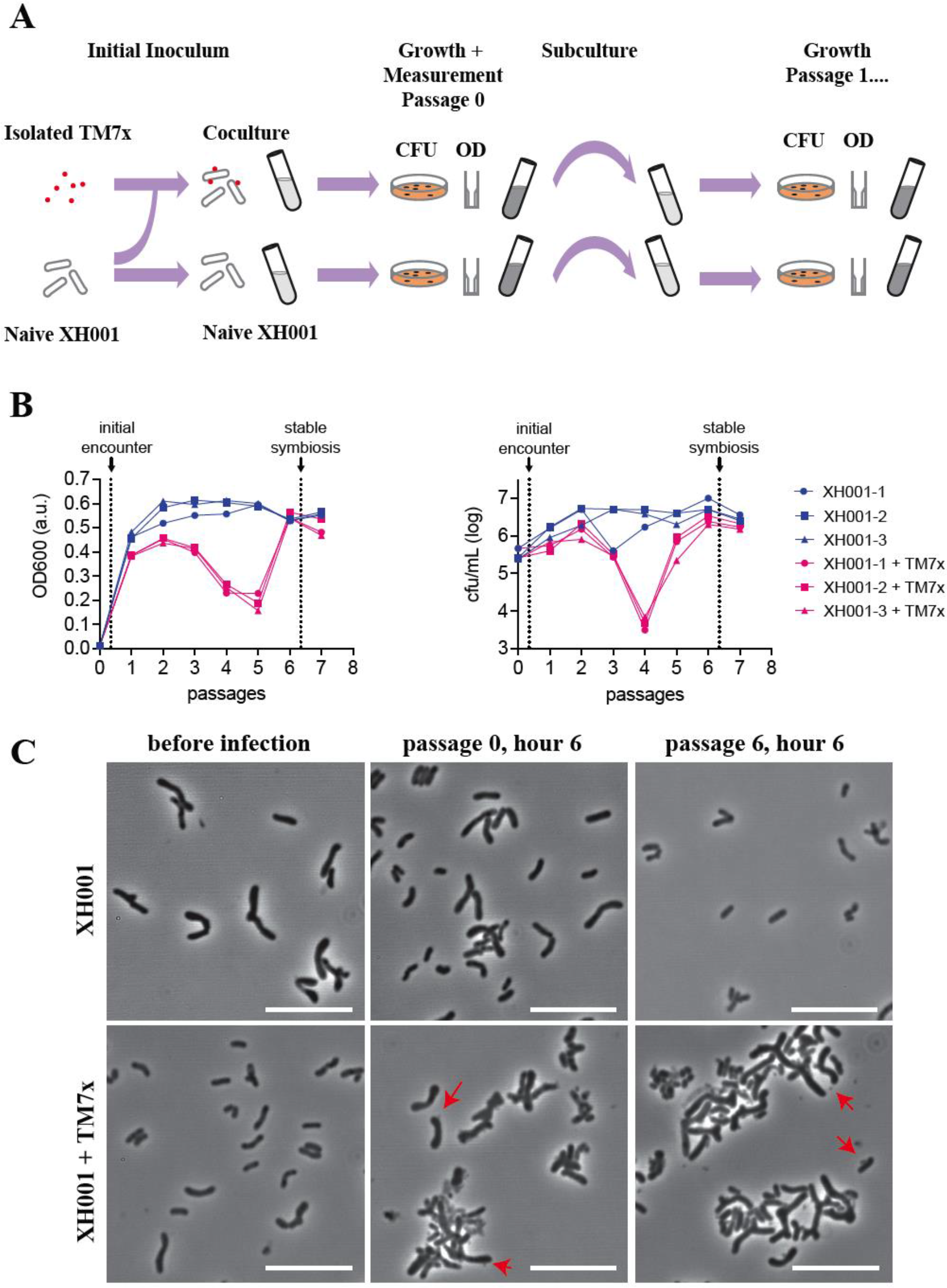
Experimental Design. A) Cartoon of culture setup, growth, and measurement for the coculture and naive cultures. Red dots are TM7x while XH001 are shown as rod shaped tubes. Optical density at 600 nm (OD600) and colony forming units (CFU) were quantified at each step of the passage and represented in panel B. B) Growth measurements for passages using both OD600 (left) and colony forming units (right) over seven passages (x-axis). Three individual XH001n cultures are shown in blue circles, squares and triangles, while cultures of XH001 with TM7x added are shown in red circles, squares and triangles. Sampling points for the initial encounter (passage 0 at 6 hours) and stable symbiosis (passage 6 at 6 hours) phases for the transcriptomics sequencing are indicated by arrows. C) Phase contrast images of XH001n or XH001/TM7x cultures at indicated sampling times are shown. TM7x infecting XH001 on the surface is indicated by red arrows. Scale bars are 10 μm.

### RNA Isolation and Sequencing

Cell pellets were resuspended in cold PBS buffer and mixed with glass beads (Ref#6914100- 98596, MPbiomedicals LLC). Using the BeadBlaster Microtube Homogenizer (BenchMark), the mixtures were shaken vigorously 3 times for 30 seconds with 30 second breaks in between at 6 m/s speed. From each sample the total RNA was isolated using the High Pure RNA isolation kit (Ref#11828665001). Isolated RNA was cleaned and concentrated with Zymo Research RNA clean & concentrator kit following kit protocol (Cat#R1015). DNA was removed using the Ambion TURBO DNA-free kit (Cat#AM1907). The RNA concentration was determined by RNA broad range Qubit kit and the nucleic acid purity was determined by Nanodrop and Qubit 3.

The total RNA was further processed by rRNA depletion with Ribozero rRNA Removal Kit (Illumina, South Plainfield, NJ, USA), and the libraries sequenced by GENEWIZ, LLC. (South Plainfield, NJ, USA) using Illumina HiSeq platform. The paired end reads were trimmed of low-quality sequence at a quality cutoff of 30 using BBduk 38.37 (14). Trimmed sequences were mapped to a reference database containing XH001 and TM7x reference sequences using Geneious prime (https://www.geneious.com) with the default settings. Mapped reads were split into XH001, with an average of 27,000,000 mapped reads per sample, and TM7x, at 880,000 reads. A subset of samples checked after sequencing for rRNA (5S, 16S, and 23S) removal success contained an average 26.5% of the raw reads of rRNA.

### Bioinformatics

Expression ratios and false discovery rates (FDR) were calculated using voom/limma (15) in Degust (16) using a significance cutoff of FDR 0.05. Annotations were assigned through consensus amongst multiple annotation platforms (1), Genbank (17), SEED (18), prokka (19), eggNOG (20), PATRIC (21), and UNIPROT (22). Blast analysis (23)was also used on selected genes. Categories of orthologous genes (COGs) were obtained from eggNOG (24). Pathways were derived from the KEGG database (25, 26).

### Cryo-ET

Exponential phase of XH001n monoculture and established XH001/TM7x coculture were prepared and cells were harvested and resuspended in PBS to a final bacterial concentration of ∼10^9 cells/mL. 100 µL of cell solution were mixed with 100 µL PBS and used for subsequent cryo-ET sample preparation.

Cryo-ET samples were prepared using copper grids with holey carbon support film (200 mesh, R2/1, Quantifoil). The grids were glow-discharged for ∼30 s before depositing 5 µl of cell solution onto them. Then, the grids were blotted with Whatman filter paper from the back side for about 8 s and rapidly frozen in liquid ethane cooled with liquid nitrogen using a home-made gravity-driven plunger apparatus.

The plunge-frozen grids were clipped into cryo-FIB AutoGrids and mounted into the specimen shuttle under liquid nitrogen. Samples were milled to thin lamellae using an Aquilos cryo-FIB system (Thermo Fisher Scientific). Samples were sputter-coated with Pt to improve the overall sample conductivity, deposited with an organometallic Pt layer (4-5 µm thick) using a gas injection system for sample protection, and milled using the gallium ion beam at 30 kV with stage tilt angle around 17°. The ion beam current was reduced according to the lamella thickness during the milling process to prevent the beam damage and finally polished with 30 pA ion beam current until ∼200 nm in thickness. Afterwards, a thin Pt layer was sputter-coated on the lamella to prevent possible charging issue during the cryo-ET imaging and the grids stored in liquid nitrogen.

The cryo-FIB lamellae were transferred to a 300 kV Titan Krios electron microscope (Thermo Fisher Scientific) equipped with a Direct Electron Detector, Volta Phase Plate (VPP) and energy filter (Gatan). SerialEM (27) was used to collect single-axis tilt series around 0 µm defocus with VPP, with a cumulative dose of ∼90 e-/Å covering tilt angles from -51° to 51° (3° tilt step). Images were acquired with an effective pixel size of 3.384 Å at the specimen level. All recorded images were first drift corrected by the software MotionCor2 (28) and then stacked by the software package IMOD (29). The tilt series were then aligned by IMOD with the patching tracking method. Tomograms were then reconstructed in IMOD using the aligned stacks.

A representative section was selected from the tomograms corresponding to different samples. The cell wall thickness was measured based on the density plot along the cell wall using IMOD (29) and ImageJ.

## RESULTS AND DISCUSSION

### Experimental design and RNA sequencing

To produce samples for RNA sequencing, free-floating TM7x cells were isolated from an established XH001/TM7x coculture. A naive XH001 strain (XH001n) was exposed to isolated TM7x at a ratio of roughly 1:1 and passaged six times. For each passage, the coculture was grown for 24 hours and then subcultured at a ratio of 1:10 into fresh media. As a control, XH001n was grown through the same passaging regime using exactly the same protocol and time scale. XH001 numbers were determined at the subculture points using both optical cell density at 600 nm (OD600) and colony forming units (CFU) (**Fig. 1A**). Due to small size, TM7x numbers have little effect on OD600 and TM7x does not form CFU without a host bacterium (12). The TM7x-exposed XH001, though not the unexposed control, underwent killing and recovery phase and by passage 6 the coculture had advanced to the stable symbiosis phase (**Fig. 1B**).

Samples for both XH001/TM7x coculture and XH001n monoculture were collected from the following passaging timepoints: 6 hours into passage 0, (termed “initial encounter phase”), and 6 hours into passage 6, (termed “stable symbiosis phase”) (**Fig 1B**). During the initial encounter, TM7x still attached to the surface of the host bacteria. Later, during the stable symbiosis phase, even more TM7x could be found on the host bacteria and the host bacteria had elongated morphology (**Fig. 1C**). For each sample, total RNA was extracted, rRNA depleted, and the libraries were sequenced using Illumina Hiseq platform (see Methods). The paired end reads were trimmed of low-quality sequence at a quality cutoff of 30. Trimmed sequences were mapped to a fused XH001 plus TM7x reference sequence. Mapped reads were split into XH001, with an average of 27,000,000 mapped reads per sample, and TM7x, at 880,000 reads. The data discussed in this publication have been deposited in NCBI’s Gene Expression Omnibus (30) and are accessible through GEO Series accession number GSE196744 (https://www.ncbi.nlm.nih.gov/geo/query/acc.cgi?acc=GSE196744).

### Functional level changes during initial encounter and stable phases

Expression ratios and false discovery rates (FDR) were calculated using voom/limma (15). Results for all genes are given in **Supplemental Table 1**. Comparisons using a significance cutoff of FDR 0.05 are shown in **Table 1** and **Figure 2**.

**Figure 2.**
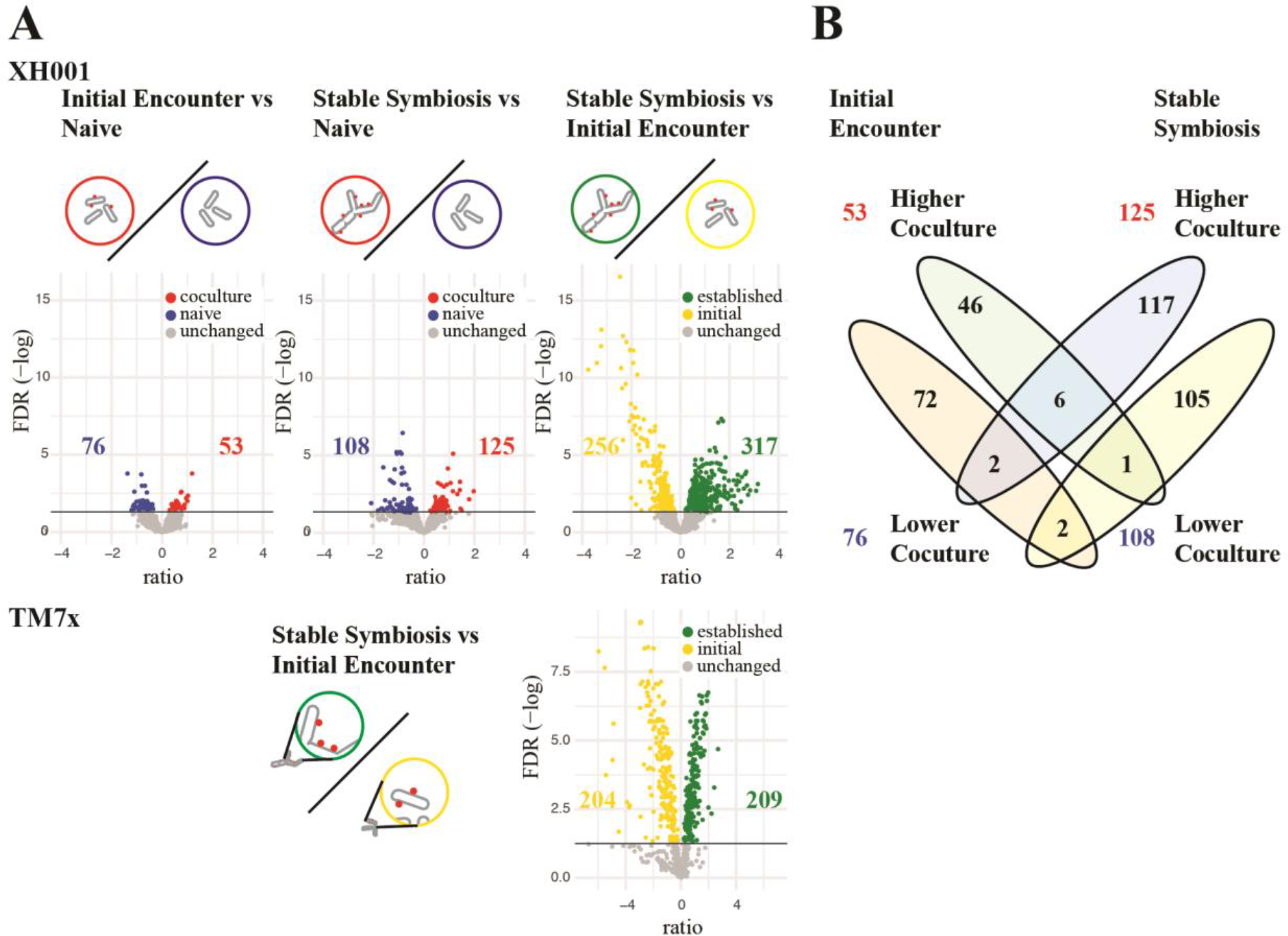
Significant Differences Between Conditions. A). Volcano plots of the log2 ratio of expression levels and -log of the FDR. Shown are all three XH001 comparisons; initial encounter coculture versus naive, stable symbiosis coculture versus naive, and stable symbiosis coculture versus initial encounter coculture. Colored dots indicate genes that made the 0.05 FDR cutoff. Blue: lower in coculture; Red: higher in coculture; Yellow: higher in initial encounter coculture; Green: higher in stable symbiosis coculture. The number of significant differences are shown on the plots. The TM7x comparison of stable symbiosis coculture versus initial encounter coculture is also shown. B). VENN diagram comparing significantly different genes for the initial encounter versus naive and stable symbiosis versus naive comparisons.

**Table 1.** Significant differences compared to controls and between phases

|  |  |  |  |
| --- | --- | --- | --- |
| **XH001 coculture**<br>**verses XH001n** | **lower with TM7x** | **unchanged** | **higher with TM7x** |
| initial encounter | 76 | 1807 | 53 |
| stable symbiosis | 125 | 1703 | 108 |
| **XH001 initial vs stable**<br>**symbiosis** | **higher in initial**<br>**phase** | **unchanged** | **higher in stable**<br>**symbiosis phase** |
| stable symbiosis vs<br>initial encounter | 256 | 1363 | 317 |
| **TM7x initial vs**<br>**stable symbiosis** | **higher in initial**<br>**phase** | **unchanged** | **higher in stable**<br>**symbiosis phase** |
| stable symbiosis vs<br>initial encounter | 204 | 279 | 209 |

Our first analysis focused on changes in host bacterium gene expression during the initial encounter. XH001n cells 6 hours after challenge with TM7x, allowing for initial binding and host response during the encounter phase, were compared to XH001n alone at the same timepoint (initial encounter versus naive control). As seen in **Table 1** and **Figure 2**, relatively few significant differences were detected for this comparison. 53 genes showed increased RNA expression in XH001/TM7x compared to the XH001n control and 76 with decreased expression. This is understandable given that XH001 and TM7x had only a short time to interact. Genes were assigned to categories of orthologous genes (COGs) using eggNOG-mapper genome-wide functional annotation as well as manual curation of annotations. As seen in **Figure 3**, the largest number of statistically significant differentially expressed genes, 30%, were in COG S: function unknown. For the other COGs, the largest groups showing increased expression in XH001/TM7x compared to XH001n were E: amino acid metabolism and transport, J: translation, G: carbohydrate metabolism and transport, and K: transcription. Decreased expression was seen in J: translation, E: amino acid metabolism and transport, P: inorganic ion metabolism and transport, and M: cell wall, membranes, and envelope biogenesis. That some of the largest numbers of increased and decreased genes were seen in amino acids metabolism and transport and translation implied a shift rather than an overall increase or decrease in these functional categories (**Fig. 3A**).

**Figure 3.**
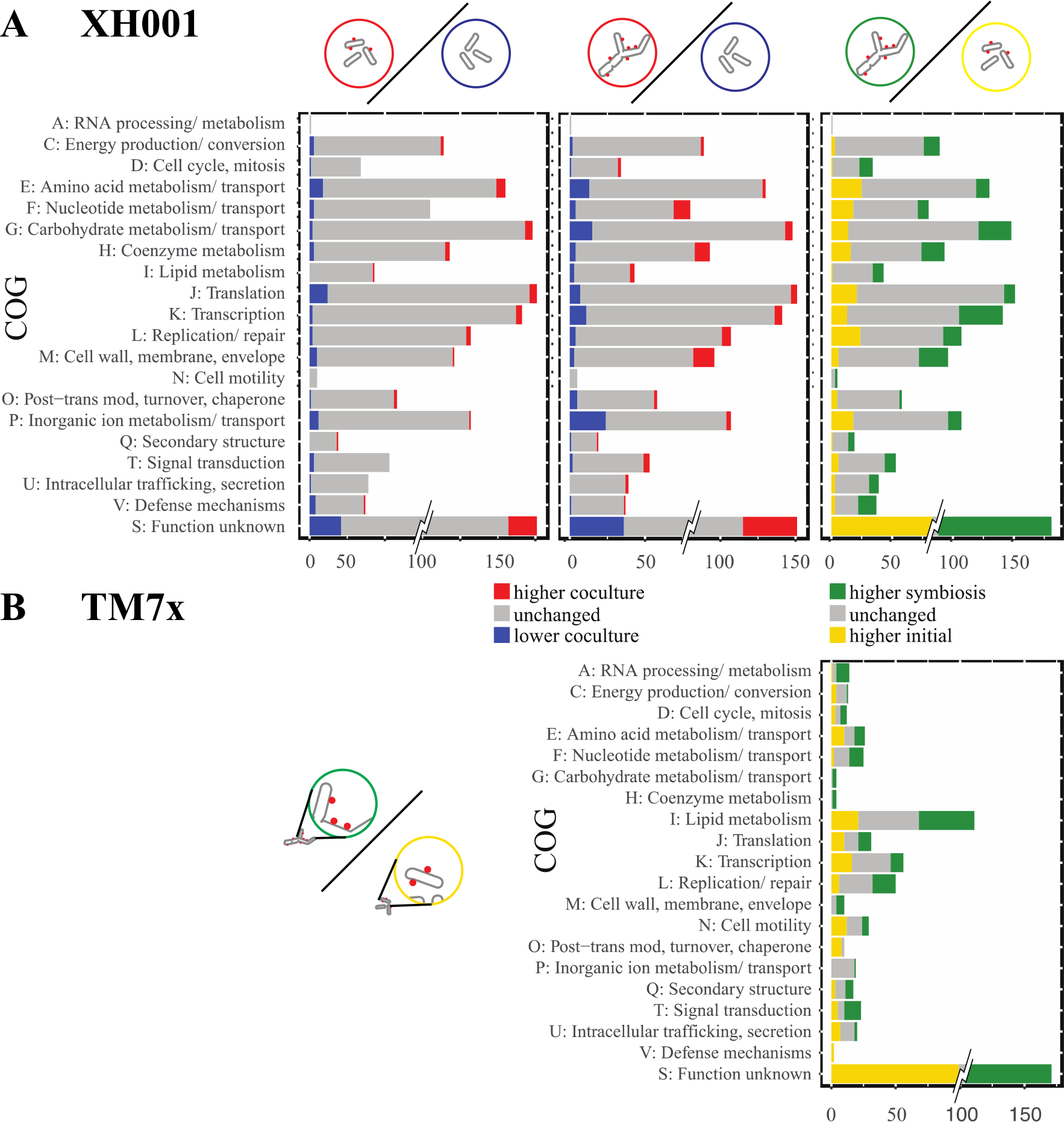
Clusters of Orthologous Groups. The number of unchanged (grey) and significantly differentially expressed genes (colored) for each COG are shown for both the XH001 (top row) and TM7x (bottom row) comparisons. To prevent the large number of genes in cluster S: unknown function from dominating the scale, only the significant differences are shown at full value. XH001 contains 621 genes annotated as S: unknown function. TM7x contains 260.

Our second analysis focused on host expression in stable symbiosis, comparing XH001/TM7x with XH001n 6 hours after subculturing into passage 6 (stable symbiosis versus naive control). This comparison detected additional significant differences between XH001/TM7x and XH001n than seen during the initial encounter, 108 increased and 125 decreased. COG analysis showed increased expression in the XH001/TM7x coculture for M: cell wall, membranes, and envelope biogenesis, F: nucleotide metabolism and transport, and H: coenzyme metabolism. Decreases were seen for P: inorganic ion metabolism and transport, G: carbohydrate metabolism and transport, E: amino acid metabolism and transport, J: translation, and K: transcription (**Fig. 3B**).

There were some consistencies between the first two comparisons. Inorganic ion metabolism and transport were decreased in the cocultures at both phases. However, COG G: carbohydrate metabolism and transport shifted from more genes with increased expression in the coculture during the initial encounter to more genes with decreased expression in the stable symbiosis. The results supported the presence of a distinctive shift between the XH001 response when first encountering TM7x and upon establishing a stable symbiosis as well as indicating that inorganic ions, amino acids, carbohydrates, and changes to the cell wall play an important role in the host/epibiont interactions.

A third analysis compared XH001/TM7x from the stable symbiosis to XH001/TM7x during the initial encounter. Significant differences were seen between these states (317 increased during stable symbiosis and 256 increased during the initial encounter) indicating a substantial shift in expression within XH001 during the change to a stable symbiosis. This result was emphasized by the limited overlap seen between the first two comparisons (**Fig. 2B**).

Most COGs showed significant differences between the initial encounter phase and the stable symbiosis phase. C: energy production and conversion, D: cell cycle, I: lipid metabolism, M: cell wall, membrane, and envelope biogenesis, and V: defense mechanisms showed predominantly more increased genes in stable symbiosis. In contrast, E: amino acid metabolism and transport, F: nucleotide metabolism and transport, J: translation, L: replication and repair, and P: inorganic ion metabolism and transport showed more increased genes during the initial encounter. These results were consistent with the idea that XH001 is providing nutrients and cell wall components to TM7x. The higher levels of defense mechanisms could imply higher stress for the parasitized host.

A fourth analysis looked at gene expression within the TM7x epibiont which have not previously been investigated. Since TM7x does not grow without a host, no monoculture TM7x control was available. TM7x RNA from the XH001/TM7x stable symbiosis was compared to the XH001/TM7x initial encounter (TM7 stable symbiosis versus initial encounter). An extensive shift was seen between the phases with a majority of genes showing significant differences. 413 out of a total of 692 genes displayed differential expression (**Table 1, Fig. 2B, Fig. 3**).

COG analysis showed that many functions had sets of genes increased in the initial encounter as well as genes increased during stable symbiosis (**Fig. 3**). However, several functions were skewed heavily towards increases in stable symbiosis. These included energy production and conversion, carbohydrate metabolism, coenzyme metabolism, lipid metabolism, intracellular trafficking and secretion, and cell motility. These pathways are mostly dependent on host substrates and correlate well with host functional changes. COGs with genes mostly increased during the initial encounter were related to cell cycle, inorganic ion metabolism and transport, and defense mechanisms.

### Cell morphology, cell cycle, and cell wall/ membrane

XH001 has shown drastic changes to cell division and cell shape when infected with TM7x (12, 31). During stable symbiosis, 48-72% of host cells were found to be associated with TM7x (12). These cells had reduced cell growth and inhibited cell division, leading to elongated and clubbed end morphology. XH001 cells also had a sticky cell surface, forming smaller aggregates in coculture (31). It was speculated that the XH001/TM7x coculture was maintained indefinitely by TM7x parasitizing new hosts from the dividing non-TM7x-associated cells (12). Using cryoEM measurements, we have found a thickening of the peptidoglycan layer from ∼15nm to more than 40nm during stable symbiosis (**Fig. 4**). Given these observations, XH001 would be expected to show changes in genes for DNA replication, cell cycle, and cell wall/ membrane biosynthesis during infection. These processes are also intimately related to each other in bacteria (32).

**Figure 4:**
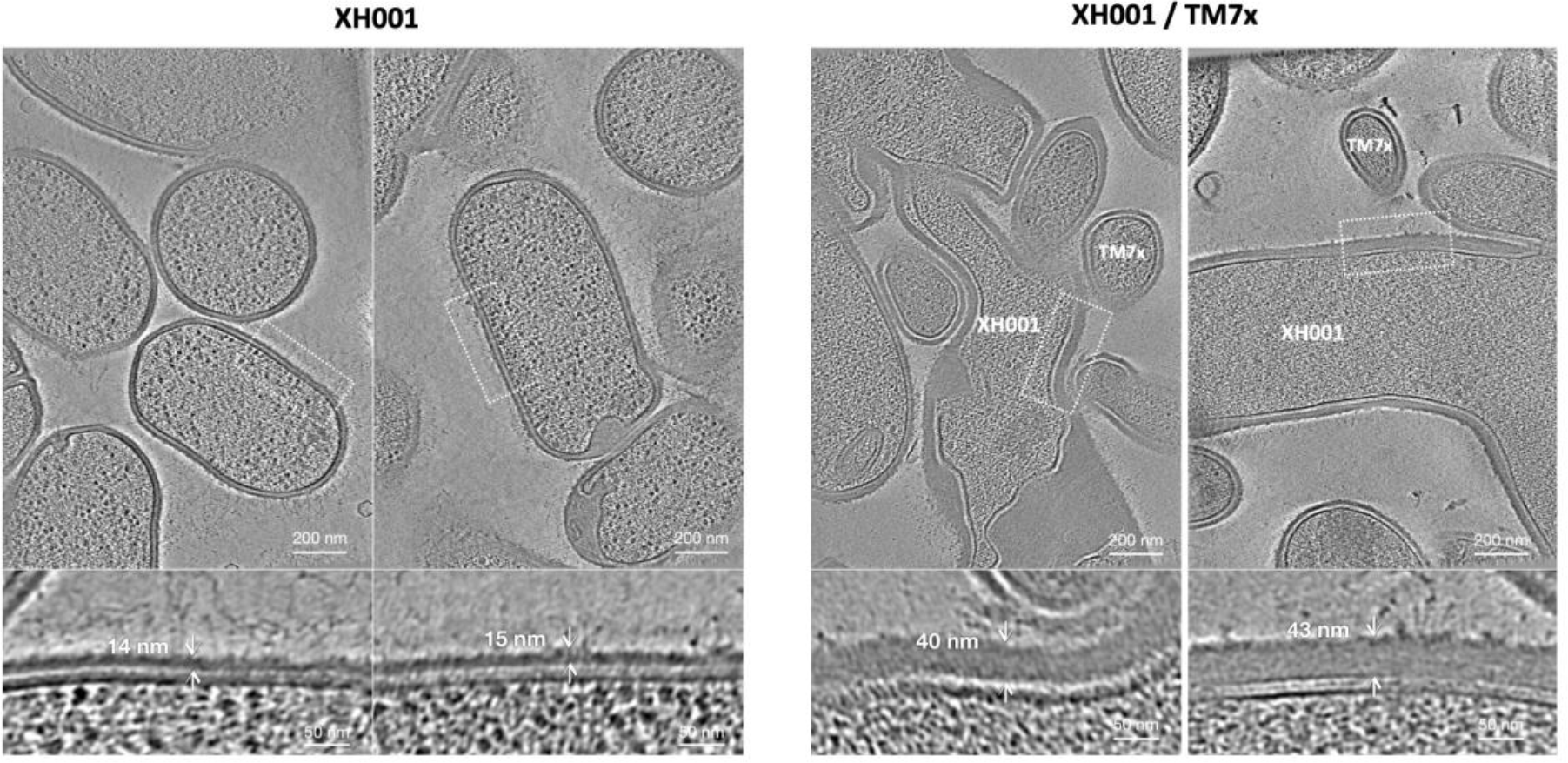
Cell wall of XH001 observed using Cryo-EM. Representative section selected from the tomograms corresponding to XH001 cells in monoculture (A), and in established XH001/TM7x symbiosis co-culture (B). The cell wall thickness was measured based on the density plot along the cell wall using IMOD and ImageJ. Top: Scale bars are 200 nm. Bottom: Scale bars are 50 nm.

The COG analysis (**Fig. 3**) indicated few differences for COG L: replication and repair genes in the first two comparisons, but a predominance of increased genes during the initial encounter when compared to stable symbiosis. Focusing on the genes for DNA replication, the first two comparisons contrasting XH001/TM7x with XH001n showed little difference for DNA replication (**Fig. 5A, Supplemental Table 2**). However, reduced RNA expression for DNA replication genes was seen during the stable symbiosis compared to initial encounter. This was consistent with the observed reduced cell growth and inhibited cell division seen in TM7x infected cells during stable symbiosis (12).

**Figure 5.**
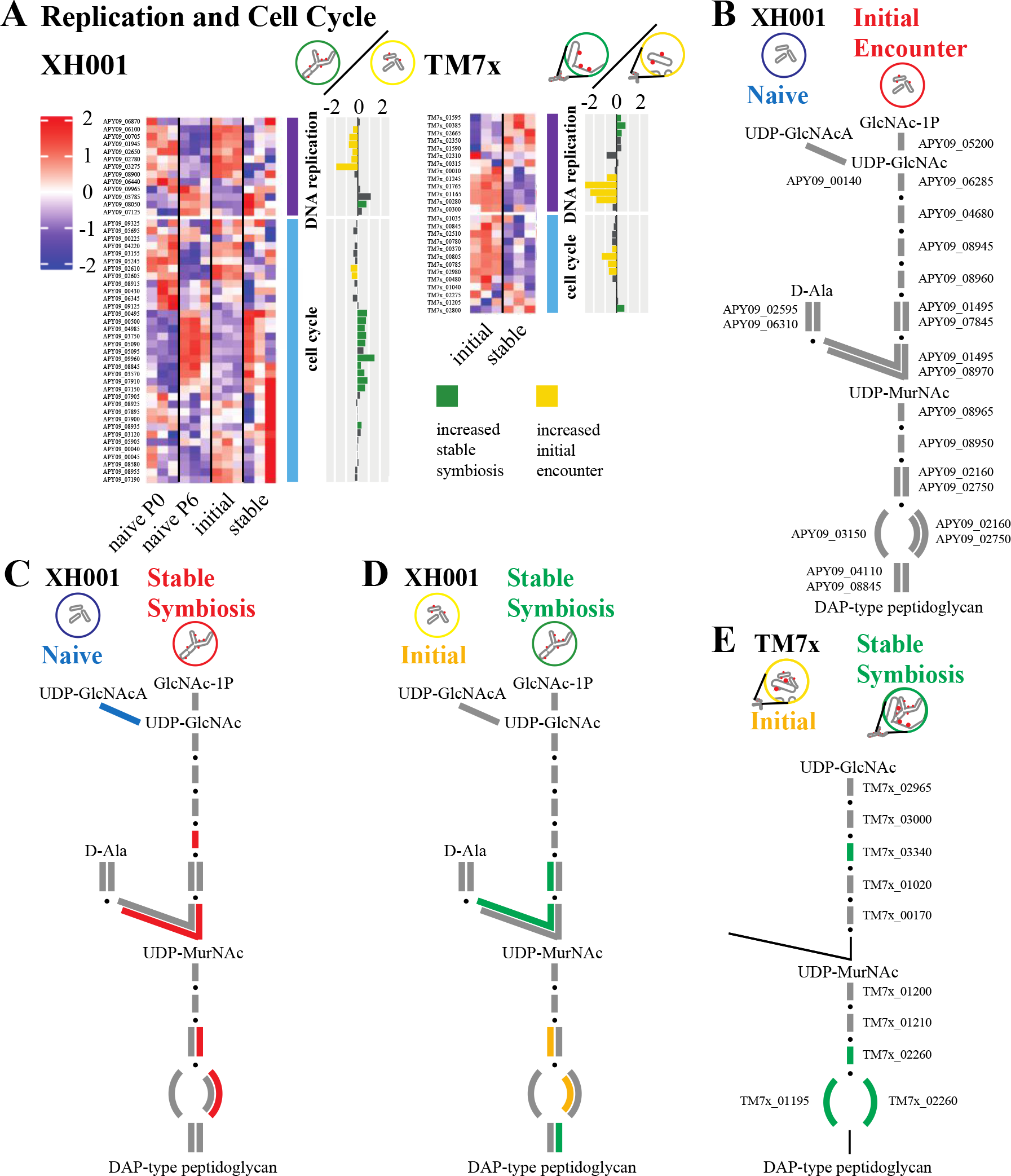
DNA replication, cell cycle, and peptidoglycan biosynthesis. A) Genes for DNA replication and cell cycle. Heatmaps of Z-score adjusted sequence counts for the three biological replicates for each condition. Bar plots show the log2 ratio between the stable symbiosis coculture and the initial encounter coculture. Green indicates significantly increased during stable symbiosis and yellow significantly increased in the initial encounter phase. B-E) A schematic of the peptidoglycan biosynthesis pathway for XH001 and TM7x. Due to its small genome TM7x has fewer predicted genes in these pathways. Steps with alternative pathways are shown as separate connections. Steps with multiple subunits are shown with multiple lines. Red: increased in XH001/TM7x versus XH001n; Blue: decreased in XH001/TM7x versus XH001n; Green: increased in stable symbiosis; Yellow: increased in initial encounter; Grey: statistically unchanged. B) Initial encounter versus naive. The XH001 APY09 gene designations are given. C) Stable symbiosis versus naive. D) Stable symbiosis versus initial encounter. E) TM7x stable symbiosis versus initial encounter. The TM7x gene designations are given. Steps that currently have no predicted gene but are expected to exist are shown in thin black lines.

The picture was more complex looking at genes for cell division control (**Fig. 5A, Supplemental Table 2**). When comparing the initial encounter to naive*, ftsK* (APY09_09125) was decreased in XH001/TM7x. However, for stable symbiosis verses naïve, two genes, including *ftsZ* (APY09_08935), were increased in XH001/TM7x, and one decreased. As seen in the COG analysis for COG D: cell cycle and mitosis (**Fig. 3**), despite DNA replication genes generally being increased in the initial encounter compared to stable symbiosis, cell cycle genes showed increased levels during stable symbiosis versus initial encounter. Out of 34 putative cell cycle genes, 11 were increased in stable symbiosis including *ftsZ* (APY09_08935) and a possible *ftsK* (APY09_09960). Interestingly, the genes significantly increased in the initial encounter were chromosome partitioning genes (APY09_02605, APY09_02610). The combination of decreased levels of DNA replication genes but increased levels of cell cycle genes during stable symbiosis compared to the initial encounter may result in the elongated and hyphal morphology seen in infected XH001(33). The lack of complete cell division would be consistent with reduced genome replication and chromosome partitioning, while genes ordinarily associated with the cell cycle might produce the elongated cell structures (33).

Over the course of infection, TM7x numbers increased on the host, reaching large numbers during the killing phase and reducing to a moderate number by the stable symbiosis phase. DNA replication and cell cycle genes in TM7x would be predicted to show increased expression in the initial encounter compared to stable symbiosis. TM7x showed mixed results for DNA replication. However, consistent with growth during the initial infection, 4 of the 13 putative cell cycle genes were increased during the initial encounter, including *ftsZ* (TM7x_00785) and *ftsK* (TM7x_00370). Only *ftsA* (TM7x_02800) was increased in stable symbiosis.

In keeping with cryoEM showing a thickened cell wall/membrane (**Fig. 4**), COG M: cell wall, membrane, and envelope biogenesis (**Fig. 3**) showed more significantly increased genes over the course of epibiont infection. XH001 has a rhamnose cell wall (1). In terms of rhamnose biosynthesis, a putative dTDP-4-dehydrorhamnose epimerase/ reductase (APY09_00110) was increased during stable symbiosis compared to XH001n (**Supplemental Table 2**). However, the only significant difference between stable symbiosis and the initial encounter encoded for the first step, glucose-1-phosphate thymidylyltransferase (APY09_00105), which had increased expression during the initial encounter.

A closer look at peptidoglycan biosynthesis genes showed no significant differences in the initial encounter versus naive comparison (**Fig. 5B**). Stable symbiosis versus naive showed three increased genes (APY09_02750, APY09_08960, APY09_08970) in the presence of TM7x (**Fig. 5C, Supplemental Table 2**), implying a possible increase in peptidoglycan production. However, stable symbiosis versus the initial encounter (**Fig. 5D**) showed mixed results, with two genes increased during stable symbiosis (APY09_01495, APY09_08845) and one increased in the initial encounter (APY09_02160). In TM7x, genes for the final synthesis steps were increased during stable symbiosis versus the initial encounter (TM7x_01195, TM7x_02260, TM7x_03340). This implied possibly higher peptidoglycan synthesis and cell division for TM7x during stable symbiosis as well.

While TM7x encodes for peptidoglycan synthesis genes, its reduced genome does not encode production of the UDP-GlcNAc building block for the pathway. Presumably the epibiont obtains UDP-GlcNAc from the host. Despite increased peptidoglycan gene RNA in XH001 and TM7x during stable symbiosis (**Fig. 5E**), XH001 did not show increases in the pathway leading to UDP-GlcNAc. However, XH001 has an alternative pathway to consume UDP-GlcNAc converting it to UDP-GlcNAcA. This gene (APY09_06140) was decreased in the stable symbiosis versus naive comparison (**Fig. 5D**). A reduction in the alternative use might increase flow of UDP- GlcNAcA into peptidoglycan synthesis.

Mannosylation, a glycosylation process found in all domains of life, can use GDP-mannose and polyprenyl phosphate mannose as mannose donors (34). As seen in **Figure 6** (**Supplemental Table 2**), instead of proceeding through glycolysis to pyruvate an alternative use for Β-D-fructose- 6P in XH001 is the production of GDP-mannose and polyprenyl phosphate mannose. There were no significant differences detected in the first two comparisons but comparing the stable symbiosis versus the initial encounter showed increased expression for genes in the pathway (APY09_05215, APY09_06140, APY09_07790) to these mannose sources. This implied increased mannosylation during stable symbiosis.

**Figure 6.**
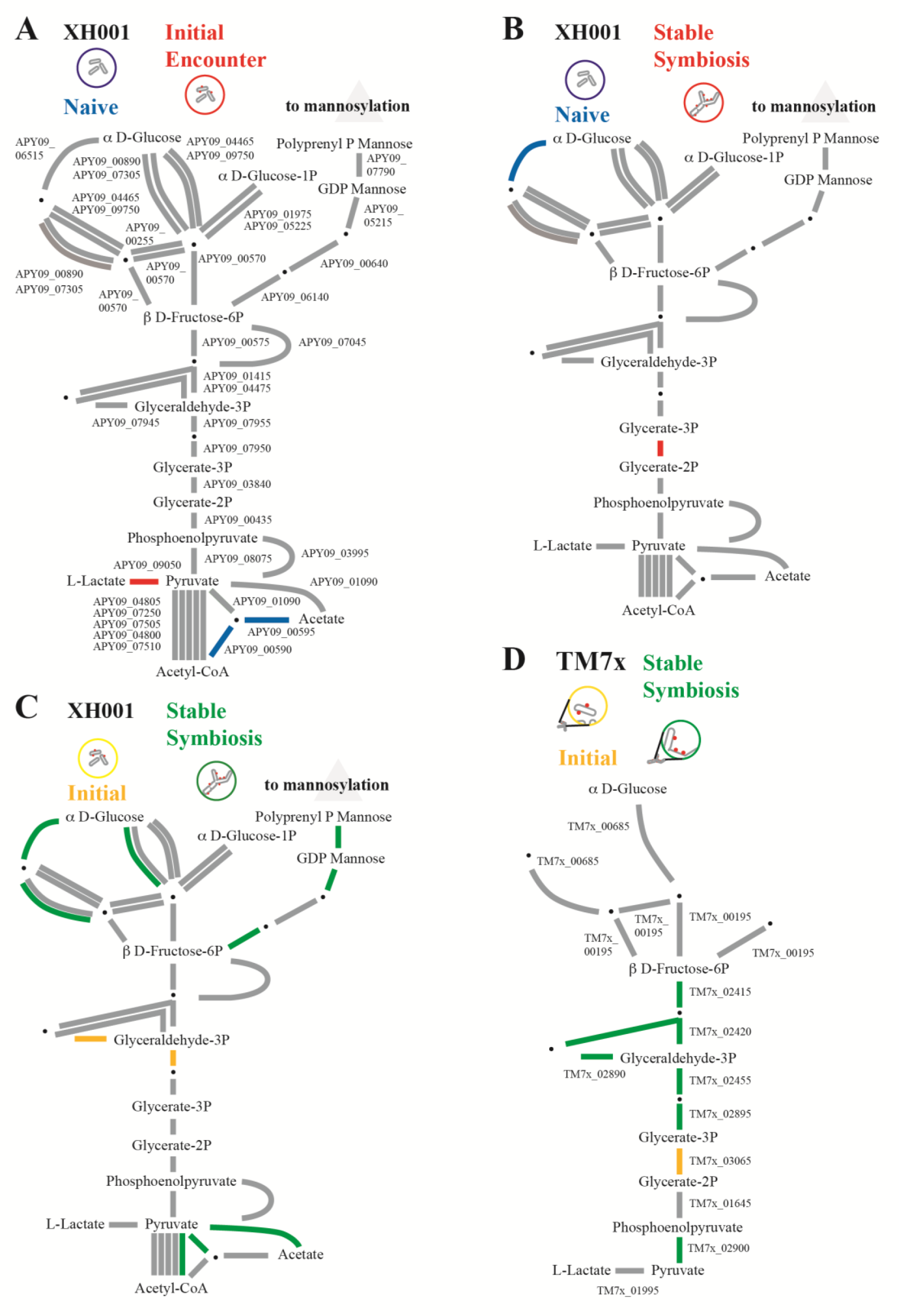
Glycolysis and Mannosylation Substrates. A schematic of the glycolysis pathway and polyprenol phosphate mannose for XH001 and TM7x. Due to its small genome TM7x has fewer predicted genes in these pathways. Steps with alternative pathways are shown as separate connections. Steps with multiple subunits are shown with multiple lines. Red: increased in XH001/TM7x versus XH001n; Blue: decreased; Green: increased in stable symbiosis; Yellow: increased in initial encounter; Grey: statistically unchanged. A) Initial encounter versus naive. The XH001 APY09 gene designations are given. B) Stable symbiosis versus naive. C) Stable symbiosis versus initial encounter. D) TM7x stable symbiosis versus initial encounter. The TM7x gene designations are given.

### Energy Metabolism

COG C: energy production and conversion indicated few differences between XH001/TM7x and XH001n in the first two comparisons but some increase in expression comparing the stable symbiosis to the initial encounter. To better understand these results we took a more detailed look at glycolysis. **Figure 6** shows the glycolysis pathway for XH001 and TM7x, though TM7x has a more limited repertoire of glycolysis genes. Comparing the initial encounter to XH001n showed few differences (**Fig. 6A, Supplemental Table 2**). However, two of the genes leading from pyruvate to acetate as an end product (APY09_00590, APY09_00595) had decreased expression while L-lactate dehydrogenase (APY09_01090) leading to l-lactate was increased in XH001/TM7x. This implied a shift in end products from acetate to l-lactate. A metabolic flux model of TM7x and the XH001 host indicated that l-lactate might play an important role in the coculture metabolism (35). A shift towards l-lactate production during the initial encounter would be consistent with the model.

Stable symbiosis versus naive control (**Fig. 5B**) also showed few significant differences. The shift from acetate to l-lactate seen in the initial encounter disappeared, possibly indicating a shift in nutrient transfer during the stable symbiosis phase. Only 2,3-bisphosphoglycerate-dependent phosphoglycerate mutase (APY09_03840) showed a significant increase in XH001/TM7x.

In contrast, comparing XH001/TM7x during stable symbiosis to the initial encounter implied lower levels for pyruvate production in symbiosis than in the initial encounter (**Fig. 6C**). Even though COG C showed more genes increased in the stable symbiosis, between B-D-fructose- 6P and pyruvate the only significant differences were triosephosphate isomerase (APY09_07945) and 6-phosphofructokinase (APY09_07955), both decreased during stable symbiosis. However, despite the decrease in these genes, stable symbiosis showed increased expression of α D-glucose- 6P (APY09_00890, APY09_06515). It is unclear why genes for the glycolysis pathway substrate would be increased during stable symbiosis while the glycolysis genes themselves would be increased during the initial encounter. As mentioned in the previously section, this might be explained by routing the higher levels of α D-glucose-6P into mannosylation during stable symbiosis rather than glycolysis.

The TM7x results comparing stable symbiosis with the initial encounter are interesting given the changes seen in the host (**Fig. 6D**). Most of the glycolysis pathway genes were increased during stable symbiosis. This is unsurprising, as TM7x would be expected to derive energy primarily from its host and this interaction should be optimal during stable symbiosis. However, 2,3-bisphosphoglycerate-dependent phosphoglycerate mutase, the only XH001 glycolysis gene increased in stable symbiosis was decreased in TM7x during stable symbiosis (TM7x_03065). One possible explanation would be nutrient exchange between the host and epibiont with TM7x effectively relying on XH001 for this step.

### Arginine

Members of the CPR are noted for their limited biosynthetic capabilities including missing pathways for de novo amino acid biosynthesis (3, 4). However, TM7x encodes for several genes in the arginine pathway that only occurs in mammalian associated members of Saccharibacteria and has been verified to utilize arginine (36). **Figure 7** shows the pathway for XH001 and TM7x. Relatively few genes in the XH001 arginine pathway showed significant differences in the first three comparisons (**Fig. 7A-C, Supplemental Table 2**), giving no strong indication of an overall change in arginine biosynthesis in the host. However, TM7x showed a significant increase in all of the genes for arginine biosynthesis/ catabolism (TM7x_03425, TM7x_03435, TM7x_03440) as well as an ornithine/arginine symporter (TM7x_03430) during stable symbiosis, implying a role for arginine in the long term epibiont interaction (**Fig. 7D**). This is consistent with the recent finding that TM7x can metabolize arginine and that it uses the arginine to stay viable and generate ATP in absence of its host bacteria (36).

**Figure 7.**
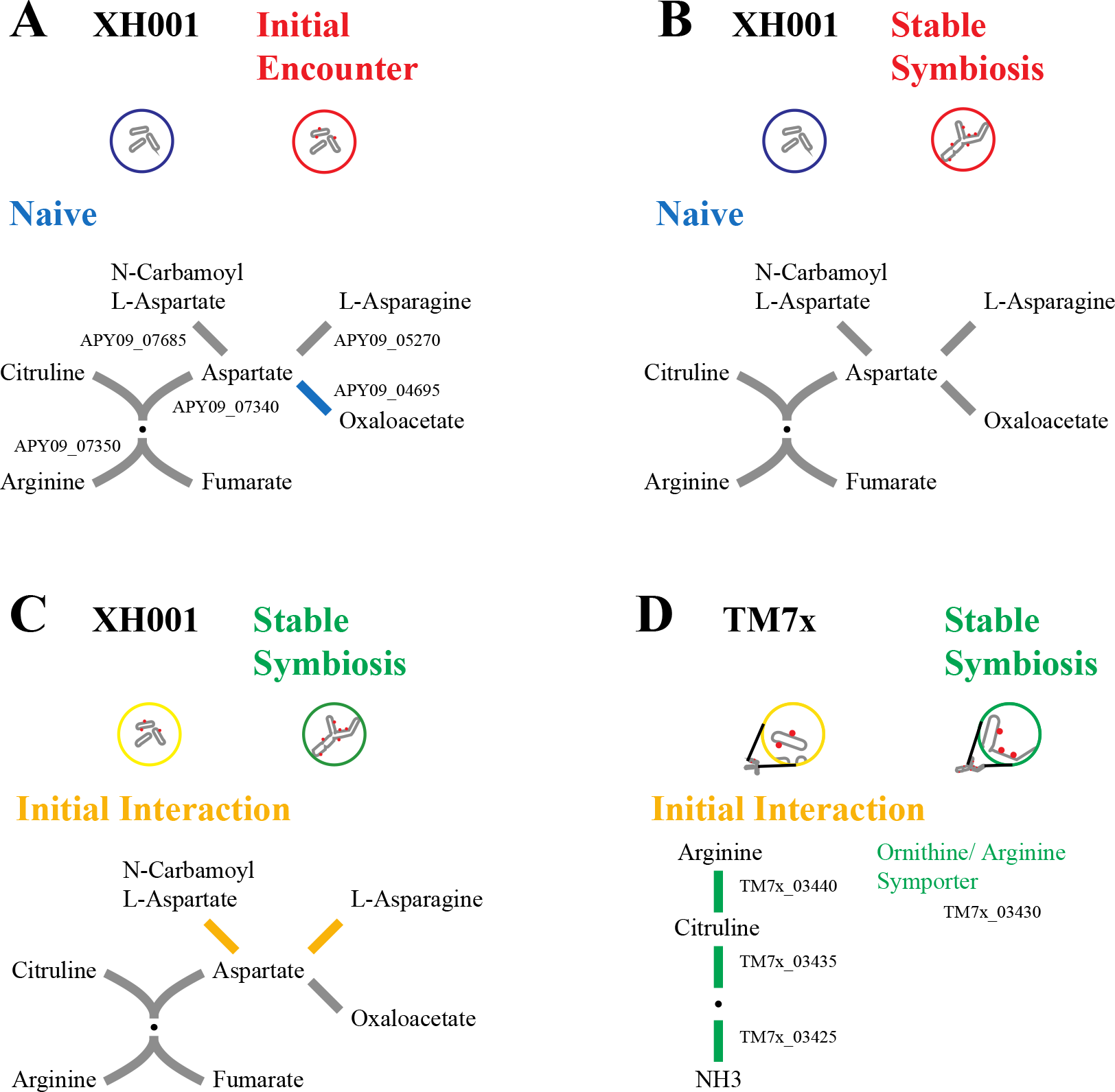
Arginine Metabolism. A schematic of the arginine pathway for XH001 and TM7x. Red: increased in XH001/TM7x versus XH001n; Blue: decreased in XH001/TM7x versus XH001n; Green: increased in stable symbiosis; Yellow: increased in initial encounter; Grey: statistically unchanged. A) Initial encounter versus naive. The XH001 APY09 gene designations are given. B) Stable symbiosis versus naive. C) Stable symbiosis versus initial encounter. D) TM7x stable symbiosis versus initial encounter. The TM7x gene designations are given.

### Autoinducer

TM7x can induce higher levels of biofilm formation in its host XH001, an effect that is dependent on autoinducer AI-2 synthase-encoding gene *luxS* and *lsrB* which encodes AI-2 transporter (37). *lsrB* (APY09_02520), showed possible differences in the first two comparisons, initial encounter versus naive and stable symbiosis versus naive, but they did not make the significance cutoff (**Supplemental Table 2**). However, *luxS* (APY09_06105) was increased in the stable symbiosis versus naive control, consistent with a role in long term associations between the species. A putative autoinducer 2 receptor-encoding gene was found in TM7x (TM7x_02705). Expression of this gene was increased in TM7x during the initial encounter than during stable symbiosis.

### Stress

Previous studies indicated that the presence of TM7x caused increased stress in XH001 (33). As seen in **Figure 8A**, the transcriptome results did not show consistent increases in stress genes for the cocultures. For the first comparison, initial encounter versus naive control, three stress related genes were increased in the presence of TM7x, RNA binding SsrA binding protein (APY09_00030), DNA repair exodeoxyribonuclease III (APY09_00470) and a putative defense involved transporter (APY09_03475) (**Supplemental Table 2**). However, five genes, heat shock chaperone (APY09_07805) and four putative transporters involved in defense (APY09_04025, APY09_04030, APY09_04035, APY09_08770), had significantly decreased expression.

**Figure 8.**
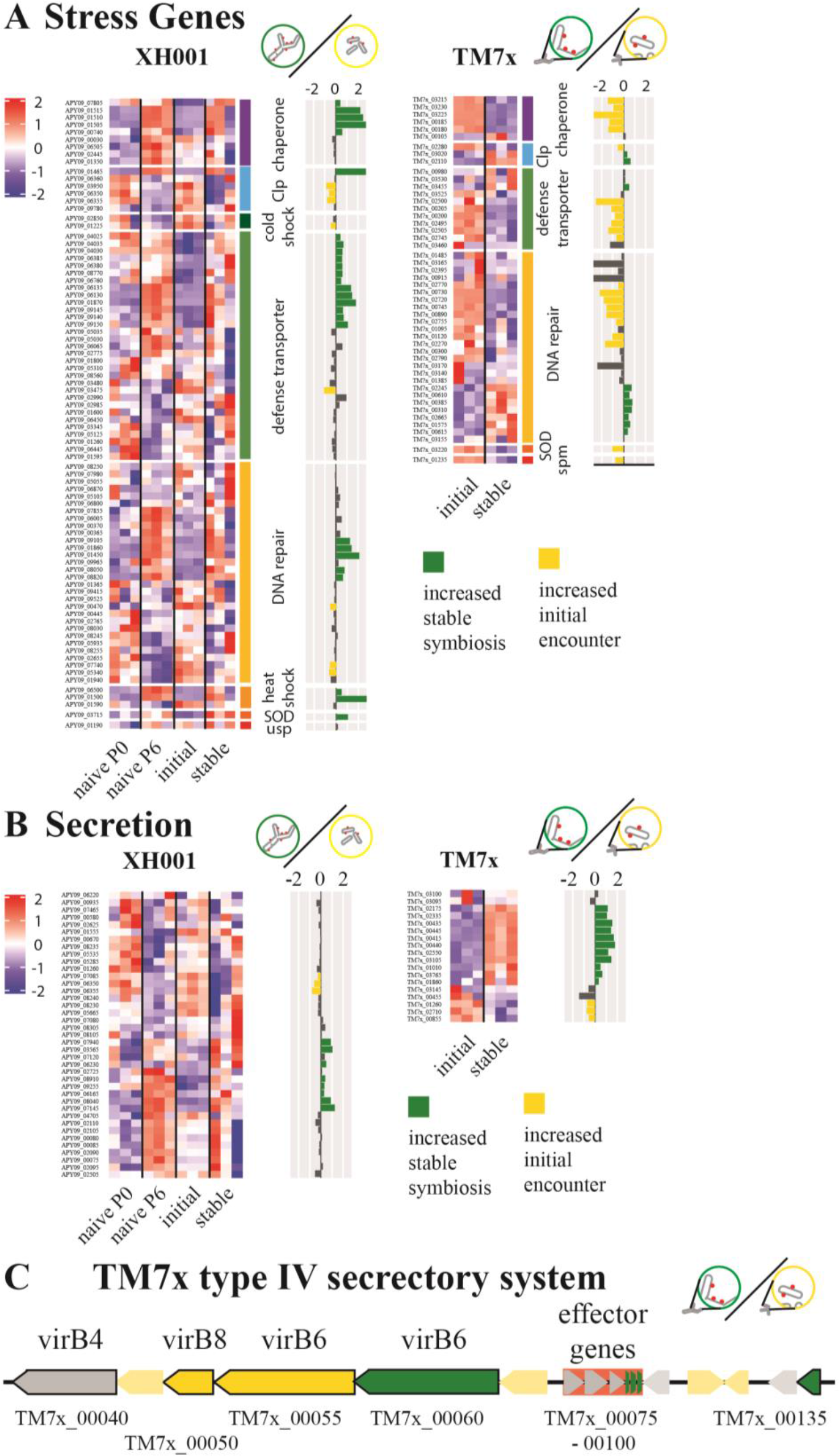
Stress and Secretory Systems. A-B) Heatmaps of Z-score adjusted sequence counts for the three biological replicates for each condition. Bar plots show the log2 ratio between the stable symbiosis coculture and the initial encounter coculture. Green indicates significantly higher during stable symbiosis and yellow significantly higher in the initial encounter phase. A) Genes involved in stress responses broken down into sub-categories chaperones, heat shock proteins, cold shock proteins, clp proteases, defense transporters, DNA repair, sodium oxide dismutase (SOD), universal stress protein (usp), and spermidine synthase (spm). B) Genes for secretory systems, excluding the type IV system from TM7x. C) A schematic of the type IV secretory system region of TM7x. Black outlines indicate predicted type IV component genes. Putative effector genes are backed in red. Green indicates significantly increased during stable symbiosis and yellow increased in the initial encounter.

The second comparison, stable symbiosis versus naive control, possibly implied reduced stress in XH001 in the presence of TM7x. Repair protein *recF* (APY09_02655) and a defense associated transporter (APY09_02990) showed increases in the coculture. However, two important chaperones *dnaJ* (APY09_06505) and *groEL* (APY09_01350) as well as a putative heat shock inducible repressor (APY09_06500) were decreased with TM7x. Additionally, DNA repair genes DNA-3-mehtyladenine glycosylase (APY09_07855) and recombination and repair protein *recO* (APY09_06005) as well as a defense transporter (APY09_05030) were also significantly decreased.

The third comparison, stable symbiosis versus initial encounter, indicated higher stress in the stable symbiosis phase. A total of 27 stress related genes had increased expression levels in the established coculture, including *clpB* (APY09_01465), *dnaK* (APY09_01515), *groES* (APY09_00740), *grpE* (APY09_01510), heat inducible repressor (APY09_06500), heat shock protein *hspR* (APY09_011500) and sodium oxide dismutase (APY09_03715). In contrast, only 8 had increased expression during the initial interaction. These included cold shock protein (APY09_01225) and both putative *clpP* (APY09_06350, APY09_06355) and *clpC* (APY09_03950) **(Fig. 8A**).

Establishing itself as an epibiont appeared to reduce stress for TM7x. In the TM7 stable symbiosis versus initial encounter comparison, 22 stress related genes were increased in the initial encounter. These included sodium oxide dismutase (TM7x_03220), spermidine synthase (TM7x_01235), and 8 genes involved in DNA repair. In contrast, only 10 genes were increased during stable symbiosis. The majority of those were also within the DNA repair group though *clpB* and *clpP* (TM7x_03020, TM7x_02110) were also increased This indicated that genes within this function are being differentially regulated and should be further investigated.

### Secretion systems and appendage genes

TM7x has minimal biosynthetic capabilities and cannot replicate without a host (1) implying that nutrient transfer is likely to play a key role in the symbiosis between host and epibiont. Host bacterial and TM7x secretion systems, therefore, could be of significant interest. COG U: intracellular trafficking and secretion (**Fig. 3**) showed few differences in the first two comparisons, but mostly increased expression during stable symbiosis compared to the initial encounter (**Fig. 8B, Supplemental Table 2**). Comparing XH001 gene expression during the stable symbiosis versus the initial encounter showed eight increased genes during stable symbiosis and three increased during the initial encounter out of thirty-nine genes with predicted secretory functions. The eight (APY09_03565, APY09_06165, APY09_06230, APY09_07145, APY09_07940, APY09_08040, APY09_08910, APY09_09255) were individual genes from different pathways including predicted allantoin permease (APY09_06230), branched chain amino acid transporter (APY09_08040), signal peptidase II (APY09_08910), and preprotein translocase subunit SecG (APY09_07940). The three increased during the initial encounter were a signal recognition particle protein (APY09_07085) and two putative ATP-dependent Clp protease proteolytic subunits (APY09_06350, APY09_06355) which have broad effects rather than focused secretory functions.

For TM7x, a majority of the predicted secretion system genes showed significant differences between stable symbiosis and the initial encounter. TM7x showed eleven genes with increased expression during stable symbiosis including a predicted signal peptidase I (TM7x_00435), preprotein translocase subunits *secA* and *secY* (TM7x_01010, TM7x_01860), a general secretion pathway protein PilB/GspE family ATPases (TM7x_02335), three pilus assembly proteins (TM7x_03105, TM7x_00445, TM7x_00415), and the twitching motility protein *pilT* (TM7x_00440). Microscopy has shown pili like structures extending from CPR members to other cells (4). The increased pilus gene expression of during stable symbiosis implied that pilus like structures may play a role in long term epibiont association. The three genes increased during the initial encounter showed some overlap with the genes increased in stable symbiosis with a predicted signal peptidase I (TM7x_00855) and preprotein translocase subunits *secG* (TM7x_02710) as well as DNA processing protein *dprA* (TM7x_01260).

Pathogens and symbionts do transfer effector molecules into hosts to modify host cell behavior, typically through a specialized apparatus such as type IV secretory systems (38). TM7x is predicted to have a type IV secretion system as well as six putative effector genes, a system that is maintained across all mammalian and environmental Saccharibacteria as well as other CPR (5, 7). **Figure 8C** shows the significant differences between stable symbiosis and the initial encounter for the type IV secretory region. Out of five predicted type IV secretion protein genes, two showed significant increases during stable symbiosis, a predicted *virB2* (TM7x_00135) and *virB6* (TM7x_00060), while two were increased during the initial encounter, another predicted *virB6* (TM7x_00055) and *virB8* (TM7x_00050). However, half of the predicted effector genes (TM7x_00090-00100) were increased during stable symbiosis, consistent with genes for altering the host cell to maintain the coculture.

### Transporters

XH001 contains 296 genes putatively annotated as transporters. A combination of consensus amongst multiple annotation platforms and blast analysis was used to group transporters by possible substrate specificity (see Methods), though many genes could only be given a general annotation as transporters. The initial encounter showed few significant differences compared to naive control (**Fig. 9A, Supplemental Table 2**). The majority of altered genes were decreased in XH001/TM7x and included peptide, iron, and some sugar transporters. Genes increased in XH001/TM7x only had general transporter predictions. This might indicate XH001 limiting transport when encountering TM7x. When comparing the stable symbiosis phase to naive control (**Fig. 9B**), the number of significantly different genes increased. There was a shift from decreased Fe transport to decreased expression for transporters of other metals.

**Figure 9.**
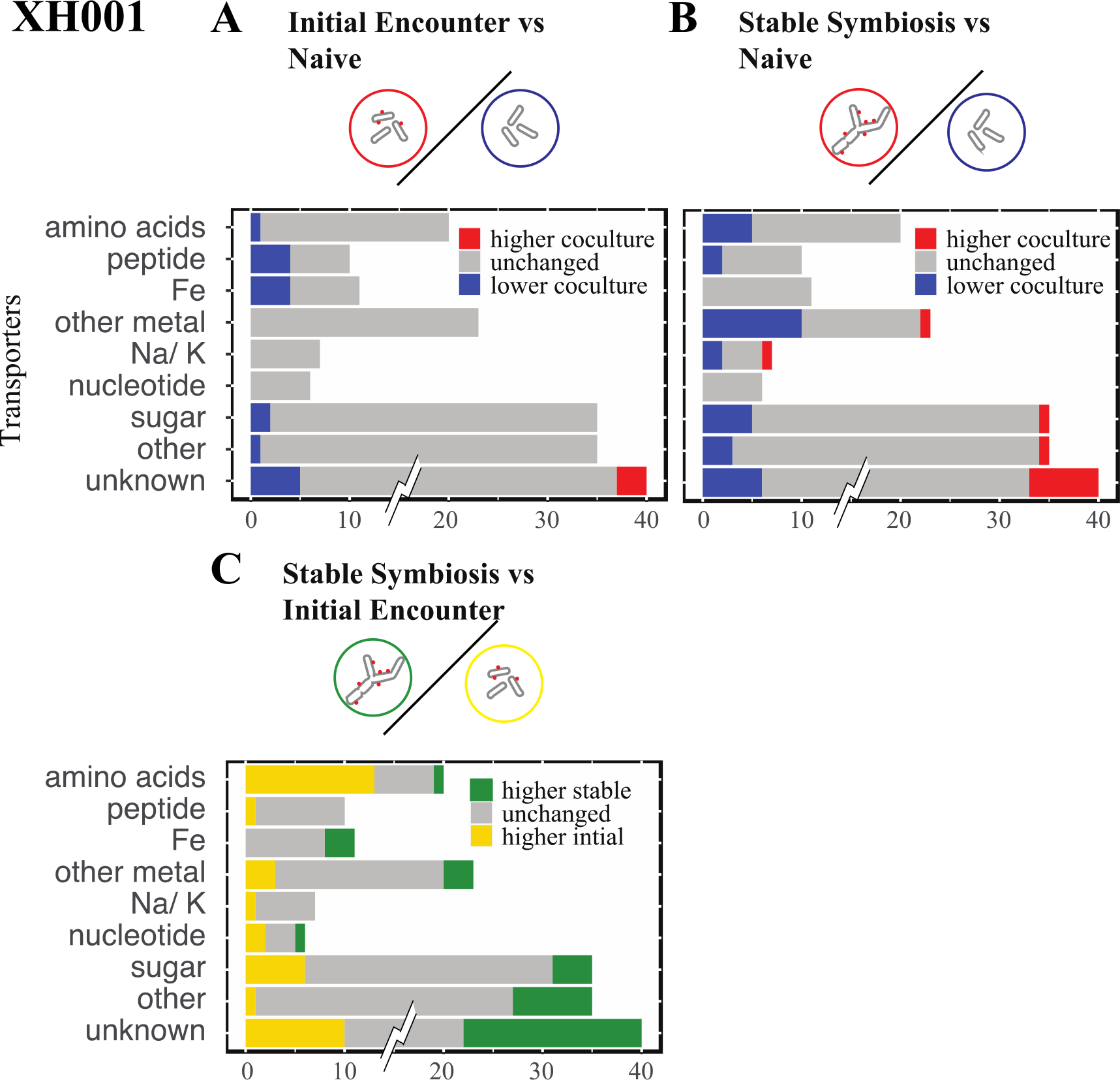
Transporters. The number of unchanged and significantly differentially expressed genes for transporter groups are shown for the XH001 comparisons. Groups: amino acids, peptides, iron (Fe), non-iron metal transporters, sodium/ potassium (Na/K), sugar, other, and genes predicted to be transporters but without an identifiable substrate (unknown). To prevent the large number of genes in unknown, 95, from dominating the scale, only the significant differences are shown at full value. A) Initial encounter versus naive control. B) Stable symbiosis versus naive control. C) Stable symbiosis versus initial encounter.

Comparing XH001/TM7x during stable symbiosis to the initial encounter reinforced the presence of a shift in expression of many transporter genes (**Fig. 9B**). A majority of amino acid transporters were increased in the initial encounter. Some non-iron metal transporters were also increased, including possible transporters of cobalt (APY09_02405, APY09_02880) and manganese (APY09_05850) transport. In contrast, iron transporters were increased during stable symbiosis. Sugar transporters showed mostly increased levels during the initial encounter though several were increased in stable symbiosis, possibly indicating a shift in sugar consumption.

TM7x has 52 putative transport genes, 23 of which showed significant increases in the initial encounter including a predicted sodium transporter (TM7x_01545), lipid A export permease (TM7x_02745), a putative copper/ silver translocating ATPase (TM7x_01845), two putative magnesium transporters (TM7x_03415, TM7x_03420), macrolide ABC transporter (TM7x_00205), multidrug ABC transporter (TM7x_02495), and a possible AI-2 receptor (TM7x_02705) discussed previously. Interestingly, there were no obvious iron transporters in the annotation. Given the importance of iron acquisition, TM7x either has a novel iron transporter that is not found in the database, or it could be acquiring iron from the host bacteria directly. We observed seventeen genes increased during stable symbiosis. Most of these were involved in secretion and were discussed in the previous section. However, these did include a putative polysaccharide permease (TM7x_03740) and a H(+) transporting ATPase (TM7x_01440). As mentioned previously an arginine/ ornithine antiporter (TM7x_03430), which could play role in host-selection (13), was also increased in the stable symbiosis phase.

## CONCLUSIONS

The CPR may constitute a quarter of bacterial diversity and its members are likely to live as epibionts on the surface of other bacteria (4), as has been conclusively demonstrated for cultivated Saccharibacteria *Nanosynbacter lyticus* type strain TM7x and its host bacterium *Schaalia odontolyticus* XH001(1) as well as within recent cultures of additional epibiont-bacterial host pairs(10, 11, 39, 40). Yet, little is understood about how these organisms establish and maintain stable symbiosis with their hosts. In prior studies, we have learned that as TM7x infects its host bacterium, there are distinct temporal phases, including an initial interaction, a killing phase, recovery phase, and finally stable symbiosis (12). To further investigate this interaction, we used RNAseq to examine both the epibiont and its host bacterium during their initial encounter and after having established a stable symbiosis. The study showed a dynamic interaction at the transcript level between the epibiont and host during the establishment of stable symbiosis.

Our data suggest that metabolites play an important role in the process. Of the few differences seen between the initial infected and naive host several centered on enzymes related to metabolites. There might have been a shift from acetate to L-lactate as an end product of energy metabolism. Additionally, the host showed lower expression of several iron, peptide, and sugar transporters. Comparing stable symbiosis to the initial encounter revealed an extensive shift in transporter expression indicating a changing interaction across the infection. TM7x also showed a significant shift in transporters between the initial encounter and stable symbiosis. Notably. the entire TM7x arginine catabolism/ biosynthesis was increased in stable symbiosis, as was an ornithine/arginine symporter. Furthermore, TM7x energy metabolism and peptidoglycan biosynthesis, which should be dependent on substrates from the host, had higher expression after establishing itself on the host.

The epibiont pili and type IV secretion system appeared to be involved in the interaction. TM7x encodes for a type IV secretion system. The secretory machinery genes did not show consistent expression changes. However, three of the putative effector genes had increased expression during stable symbiosis, implying a role for the effectors in the long-term association of epibiont and host. Microscopy has also shown pili in ultrasmall cells from fractions enriched in CPR genomes possibly interacting with hosts (6). Consistent with this, several pili genes in this study showed increased expression during stable symbiosis.

Stable symbiosis has been shown to have a dramatic effect on host cell shape, resulting in the elongated and hyphal morphology (12). We have also shown a very distinct phenotype in infected hosts with thickening of the cell wall (**Fig. 4**). Consistent with this extensive cellular change, numerous XH001 genes involved in cell wall and membrane biosynthesis were increased during stable symbiosis. Cell cycle genes also showed higher expression, with the exception of reduced expression of the chromosome partitioning genes.

The interaction also appeared to be stressful to the host. A previous study showed increased stress gene expression in the presence of TM7x (33) . Additionally, host cells showed physiological changes, growth arrest and elongated cell shape (12). Consistent with these results, most of the XH001 stress related genes showed increased expression during stable symbiosis. In contrast, the majority of epibiont TM7x stress genes were decreased after establishing itself on the host. Overall, this study has revealed the gene expression of the host bacterium and epibiont during this dynamic interaction which represents initial insights into the mechanisms of how these unique microbes, with limited de novo biosynthetic capabilities, may survive and persist at high prevalence within the human microbiome as well as in the environment.

## ACKNOWLEDGMENTS

This research was supported by grants from the National Institute of Dental and Craniofacial Research of the National Institutes of Health under Awards 1K99DE027719–01 (BB); T90DE021984 (KAK); 1R01DE023810 (XH, WS and JSM).

